# *In vivo* evaluation of the effect of sickle cell hemoglobin S, C and therapeutic transfusion on erythrocyte metabolism and cardiorenal dysfunction

**DOI:** 10.1101/2023.02.13.528368

**Authors:** Angelo D’Alessandro, S. Mehdi Nouraie, Yingze Zhang, Francesca Cendali, Fabia Gamboni, Julie A. Reisz, Xu Zhang, Kyle W. Bartsch, Matthew D. Galbraith, Victor R. Gordeuk, Mark T Gladwin

## Abstract

Despite a wealth of exploratory plasma metabolomics studies in sickle cell disease (SCD), no study to date has evaluate a large and well phenotyped cohort to compare the primary erythrocyte metabolome of hemoglobin SS, SC and transfused AA red blood cells (RBCs) *in vivo*. The current study evaluates the RBC metabolome of 587 subjects with sickle cell sickle cell disease (SCD) from the WALK-PHaSST clinical cohort. The set includes hemoglobin SS, hemoglobin SC SCD patients, with variable levels of HbA related to RBC transfusion events, and HbF related to hydroxyurea therapy. Here we explore the modulating effects of genotype, age, sex, severity of hemolysis, and hydroxyurea and transfusion therapy on sickle RBC metabolism. Data - collated in an online portal – show that the Hb SS genotype is associated with significant alterations of RBC acylcarnitines, pyruvate, sphingosine 1-phosphate, creatinine, kynurenine and urate metabolism. Surprisingly, the RBC metabolism of SC RBCs is dramatically different from SS, with all glycolytic intermediates significantly elevated in SS RBCs, with the exception of pyruvate. This result suggests a metabolic blockade at the ATP-generating phosphoenolpyruvate to pyruvate step of glycolysis, which is catalyzed by redox-sensitive pyruvate kinase. Increasing in vivo concentrations of HbA improved glycolytic flux and normalized the HbS erythrocyte metabolome. An unexpectedly limited metabolic effect of hydroxyurea and HbF was observed, possibly related to the modest induction of HbF in this cohort. The metabolic signature of HbS RBCs correlated with the degree of steady state hemolytic anemia, cardiovascular and renal dysfunction and mortality.

**Key points:**

- In vivo dysregulation of RBC metabolism by HbS is evaluated by metabolic profiling of 587 patients with variable HbA, HbC and HbF levels;
- RBC acyl-carnitines, urate, pyruvate metabolism, S1P, kynurenine relate to hemolysis and cardiorenal dysfunction, respond to transfusion;

## INTRODUCTION

Owing to its critical role in oxygen transport, hemoglobin is by far the most abundant protein in mature red blood cells (RBCs – 250-270 million copies per cell),^1^ the most abundant cells in the human body (∼83% of total host cells).^2^ At different stages of life — embryonic, fetal and adult – different genes encode for multiple types of globin proteins; as such, various tetrameric combinations of globin chains generate multiple types of hemoglobin (Hb): Hb A (HbA) is the most abundant (>90%) form of adult Hb, comprising two α-globin subunits (encoded by the duplicated *HBA1* and *HBA2* genes) and two β-globin subunits (encoded by *HBB*). Sickle cell disease (SCD) is a group of inherited disorders caused by mutations in the *HBB* gene.^3^ Single nucleotide substitutions in *HBB* result in sickle Hb (HbS) allele β^S^, owing to the substitution of amino acid E6V, or in HbC, E6K substitution. Deoxygenation of mutant globins induces polymerization, altering erythrocyte structure, function, rheology and metabolism, and represents the primary upstream mechanism for this complex disease, characterized clinically by hemolytic anemia, inflammation and recurrent vaso-occlusive episodes.^4^ In children, SCD may manifest with the development of cerebrovascular disease and cognitive impairment, requiring lifelong transfusions.^4^ As SCD subjects age, recurrent episodes of vaso-occlusion and inflammation result in progressive damage to most organs, including the brain, kidneys, lungs, bones, and cardiovascular system.^4^

High-energy phosphates adenosine triphosphate (ATP) and 2,3-diphosphoglycerate (DPG) promote off-loading of oxygen from hemoglobin through stabilization of the tense, deoxygenated tetramer.^5^ While beneficial in healthy subjects facing hypoxia, in SCD these compounds promote HbS and C polymerization, aggravating disease severity. Recent application of single cell oxygen kinetics measurements and metabolomics^6^ has identified novel small molecule candidates that may indirectly contribute to oxygen kinetics. For example, the lipid sphingosine 1-phosphate (S1P), which is synthesized by RBCs in response to hypoxia,^7^ promotes glycolysis, DPG synthesis and Hb-S polymerization in murine models of SCD and human SCD RBCs.^8^ Overall, modulation of RBC glycolysis and the Rapoport-Luebering shunt, the pathways through which ATP and DPG are synthesized in the mature RBC, seem to participate in the modulation of clinical sequelae to SCD, such as renal and pulmonary dysfunction.^9-11^ The primacy of RBC metabolism in the regulation of SCD sequalae is further illustrated by the promising preliminary evidence on the efficacy of drugs that promote DPG consumption by activating late glycolytic enzyme pyruvate kinase (PK).^12^ Similarly, common polymorphisms that affect rate-limiting enzymes of antioxidant pathways can compound with SCD by increasing susceptibility to oxidant stress-induced hemolysis.^13-15^ Such is the case of glucose 6-phosphate dehydrogenase deficiency, a condition that – like sickle cell traits - underwent positive evolutionary selection in areas where malaria is endemic, and negatively impacts RBC antioxidant metabolism through ablation of the pentose phosphate pathway (PPP).^16,17^

Application of omics technologies, particularly metabolomics, in the context of SCD has identified additional mechanisms of morbidity in the SCD population. For example, dysregulation of arginine metabolism^18^ may constrain nitric oxide biosynthesis through substrate competition of arginase with endothelial nitric oxide synthase, thus limiting the vasodilatory capacity in SCD, with direct upstream regulatory effects on vaso-occlusive crisis, cardiovascular, cerebral and renal complications of SCD.^19-24^ While most metabolomics studies in SCD have focused on plasma, a limited set of studies has investigated the RBC compartment.^11,20,22,25,26^ These studies - always limited in sample size owing to technical constraints - suggested that depletion of glutamine, a precursor to glutathione, is a potential contributor to the increased susceptibility to hemolysis of the sickle erythrocyte,^26^ owing to oxidant damage to proteins and lipids.^27,28^ The Lands cycle^27^ is the main pathway through which the mature RBC, which cannot synthesize de novo long chain fatty acids and complex lipids, repairs damaged components at the membrane level by leveraging acylcarnitine pools to preserve RBC membrane deformability.^29^ Unchecked membrane lipid damage primes extravascular hemolysis^30^ through splenic sequestration and erythrophagocytosis by the reticuloendothelial system.^31^

The implementation of high-throughput metabolomics has fueled the promise of omics applications in personalized medicine in the field of SCD and allows for the identification of upstream metabolic targets for new therapy.^32^ However, most studies have hitherto leveraged pharmacogenomics, and no metabolomics study to date has tested enough subjects with biological, genetic and clinical heterogeneity to investigate the impact of factors such as genotype, age, sex, and modulating effects of HbA and and HbF on the metabolome of SCD. To bridge this gap, here we leveraged the WALK-PHaSST cohort to perform the largest RBC metabolomics study on SCD to date. Omics data were mined in light of critical biological, clinical and genomics covariates available as part of this well-curated cohort.^23,33-37^

## METHODS

An extended version of this section is provided in **Supplementary Methods Extended**.

### Ethical statement

All experimental protocols were approved by the University of Pittsburgh, as part of the WALK-PHaSST clinical trial (NCT00492531 at ClinicalTrials.gov and HLB01371616a BioLINCC study).

### Blood collection, processing and storage

Subject enrolling, eligibility and exclusion criteria were extensively described in prior publications.^23,34-38^ In summary SCD patients were recruited in Treatment of Pulmonary Hypertension and Sickle Cell Disease With Sildenafil Therapy (walk-PHaSST) study screening phase. A total of 587 SCD patients with available RBC samples were used for metabolomics analyses. Clinical covariates were collected prior to metabolomics analysis, including hemolysis, tricuspid regurgitant velocity (TRV),^39^ parameters indicative of kidney function such as estimated glomerular filtration rate (eGFR), creatinine, and percentage of hemoglobin S (sickle - HbS), C (HbC), F (fetal - HbF) and A (alpha, normal - HbA). Description of these measurements had been previously described.^23,34-38^ Samples were then stored at -80LJ until further processing.

### Ultra-High-Pressure Liquid Chromatography-Mass Spectrometry (MS) metabolomics

Metabolomics extractions,^40,41^ analyses^42,43^ and elaborations^44^ were performed as previously described.

### Statistical analyses

Multivariate analyses, including principal component analyses (PCA), hierarchical clustering analyses, two-way ANOVAs, correlation analyses (Spearman) and calculation of receiver operating characteristic (ROC) curves were performed using MetaboAnalyst 5.0.^45^ For survival analysis, we used time to right censorship (including death or last follow-up) as time to event and vital status (death or alive) was the studied outcome. PCA was used to derive a hemolytic component from lactate dehydrogenase, aspartate aminotransferase total bilirubin and reticulocyte percent.^14,35,46^ We applied Cox proportional hazard models to calculate the hazard ratios and P value for each metabolite. and Cox analysis and adjusted regressions of metabolites to clinical outcomes were performed in R (R Core Team (2022), https://www.r-project.org/).

### Sickle Cell Disease ShinyApp porta

A portal for online sharing of all the data generated in this study, based on code from the COVID-Ome Explorer portal.^47^

### Data sharing statement

All results described in this study are provided in **Supplementary Table 1** and accessible at https://mirages.shinyapps.io/SickleCellDisease/,

## RESULTS

### Metabolomics of the WALK-PHASST SCD cohort

Results are reported in tabulated form in **Supplementary Table 1**, along with subject (n=587) biological data, genotype, sex, age, body mass index (BMI – **Figure 1.A**). Hemoglobin profiles, including percentages of hemoglobin S (HbS), A (HbA), C (HbC) and F (HbF) were measured and used along with genotyping to classify the main groups of interest, and attempt to characterize the unique effect of intracellular HbS on in vivo RBC metabolism.

**Figure 1.**
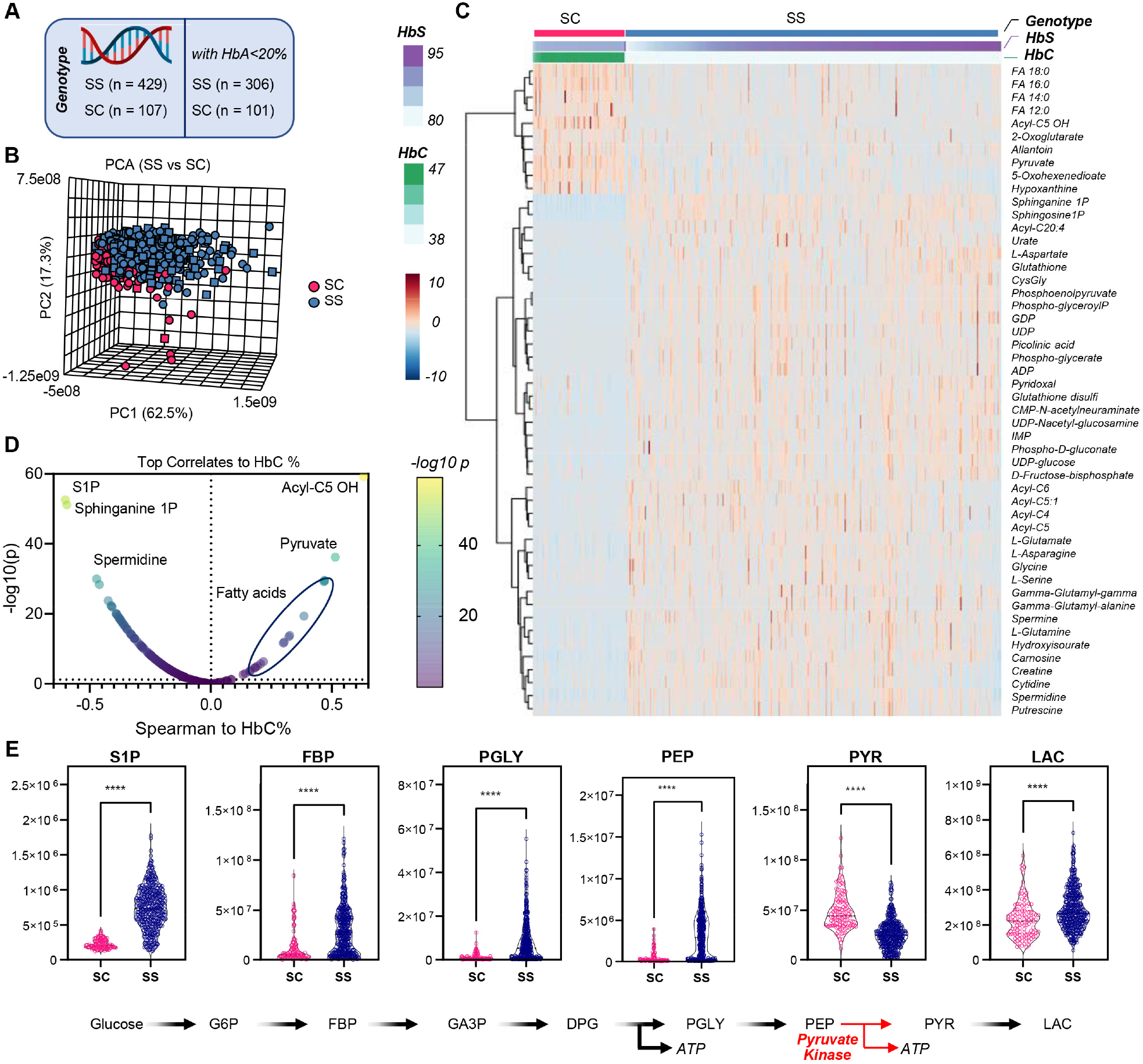
RBC metabolic differences between SCD patients with SS and SC genotypes. RBC metabolism of patients with SS (n=306) or SC genotype (n =107 - **A**) who were not recently transfused (HbA <20%) were compared via principal component analysis (**B**), hierarchical clustering analysis of the top 50 metabolites by t-test (**C**), Spearman correlation analyses between metabolites and HbC % (**D**). In **E**, focus on selected metabolites, including sphingosine 1-phosphate (S1P) and glycolytic metabolites between SC and SS RBCs.

### RBC metabolic profiles in SS vs SC patients who have not been recently transfused

First, we focused on individuals with the SS genotype (n=429) as opposed to SC genotypes (n=107), the latter characterized by reduced levels of steady state hemolytic anemia.^48^ For this analysis we aimed to limit the bias of recent transfusions, with exclusion of patients with HbA% >20%, resulting in a total of 306 SS and 101 SC samples (**Figure 1.A**). PCAs show partially separated clustering of SS and SC samples along PC2, explaining 17.3% of the total variance (**Figure 1.B**). Hierarchical clustering analysis (HCA) of the top 50 metabolites by p-value highlighted higher levels of saturated free fatty acids (C12, 14, 16, 18 series), carboxylates (2-oxoglutarate, pyruvate) and hydroxyisovalerylcarnitine (Acyl-C5 OH) derived from branched chain amino acid metabolism in the SC group (**Figure 1.C**). SS genotype was associated with elevation in the levels of S1P, nucleoside mono- and diphosphate (ADP, GDP, UDP, IMP), oxidized glutathione (GSSG) and multiple byproducts of the gamma-glutamyl cycle, purine oxidation products (urate), amino acids (asparagine, aspartate, glycine, glutamine, serine), neurotoxic kynurenine metabolites (picolinic acid) and polyamines (spermidine, putrescine – **Figure 1.C**). Linear correlation of metabolomics data to HbC% showed a strong negative correlation with S1P and its sphinganine metabolite, and a strong positive association with pyruvate and multiple fatty acids (**Figure 1.D**). This observation is interesting in that all glycolytic intermediates were significantly elevated in SS RBCs, with the exception of pyruvate, suggesting a metabolic blockade at the ATP-generating phosphoenolpyruvate to pyruvate step of glycolysis which is catalyzed by PK (**Figure 1.E**). These results were recapitulated both upon exclusion of any SS patients who received any transfusion at all (inclusion criterion: HbA 0% - **Supplementary Figure 1**) or upon inclusion of all patients, independently from transfusion status and without adjustment for any relevant covariates such as age (**Supplementary Figure 2**). Similarly, elevation in glutamine, glutamate, aspartate, but lower 2-oxoglutarate; elevation of urate and S1P, as well as arginine metabolite ornithine were observed across all SS subjects whencompared to SC patients, with elevation of citrulline and creatinine positively associated with the age of the subject (**Supplementary Figure 3**).

### Impact of transfusion on RBC metabolism in SS patients

To estimate the therapeutic effect of RBC transfusion and further characterize the effect of HbS on RBC metabolism in vivo, we analyzed the metabolite association relative to the % of hemoglobin that was HbS vs HbA in subjects who were not receiving hydroxyurea (n=243). HCA of the 50 most significant metabolites by HbS and HbA was performed by sorting results based on HbS and HbA% (**Figure 2.B –** data adjusted for relevant covariates, such as subject sex, age, body mass index). Expectedly, mannitol - a component of most additive solutions that are used for storage of packed RBCs - was identified as a top covariate to HbA% (**Supplementary Figure 4)**, when including all SS patients independently from other variables (n=429). Compared to Hb SS RBCs, transfused Hb AA RBCs were associated with higher levels of reduced glutathione (GSH) and other antioxidants (ascorbate, carnosine), early steps-glycolytic metabolites (glucose, fructose bisphosphate), PPP metabolites (6-phosphogluconate), purine monophosphates (AMP, GMP, IMP), and certain amino acids or monoamines (aspartate, dopamine, adrenaline - **Figure 2.B**). Vice versa, compared to transfused Hb AA RBCs, Hb SS RBCs were associated with elevation in the whole pool of acylcarnitines (C6, 8, 10:0, 10:1, 12:0, 14:0, 14:1, 16:0, 16:1, 18:0, 18:1, 18:2), creatine and phosphocreatine, carboxylic acids (2-hydroxyglutarate, malate), purine, methionine and glutathione oxidation products (hypoxanthine, methionine *S*-oxide, 5-oxoproline), short and medium-chain fatty acids (butanoate, pentanoate, octanoate) and a different group of amino acids (alanine, asparagine, glycine, serine, threonine - **Figure 2.B**) To further define the effect of Hb A on SS metabolism, we focused on SS subjects with extreme HbA% - i.e., lower than 5% (not recently transfused – n=147) versus HbA% >80% (recently undergoing exchange transfusion – n=19 – volcano plot in **Figure 2.C**). This analysis made the above changes even more evident, with the additional observation of elevated 2-oxoglutarate, linoleic acid (FA 18:2) and decreased lactoylglutathione and indole-pyruvate (tryptophan metabolite of bacterial origin) in high HbS% SS patients (**Figure 2.C**). Results were further supported by Spearman correlations of metabolites against HbS% (**Figure 1.D**). In summary, we identified five main pathways that differ between high HbS and high HbA groups including elevated purine catabolism, PPP, glycolysis, lipid remodeling and Lands cycle (**Figure 2.E)**.

**Figure 2.**
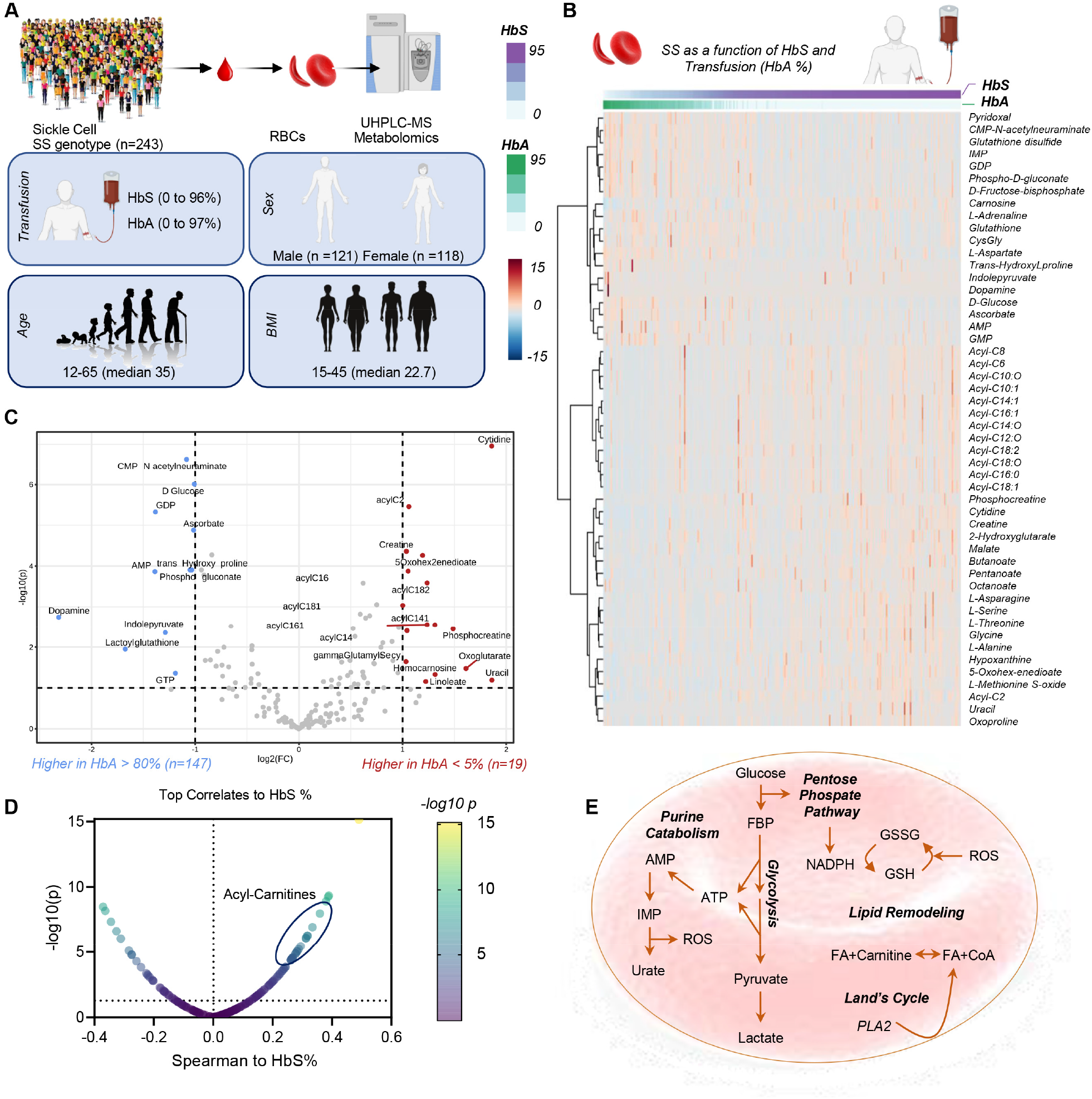
Metabolomics of the WALK-PHASST SCD cohort. Patients with SS genotype – who did not receive other treatment (e.g., hydroxyurea) were evaluated in the function of recent transfusion events, as determined by HbA % (n=243 - **A**). In **B**, hierarchical clustering analysis of metabolomics data as a function of HbS and HbA% (top 50 significant metabolites are shown). In **C**, volcano plot of metabolites significantly higher in patients with no recent transfusion event (HbA <5%) or with recent exchange transfusion (HbA>80%). In **D**, smile plot of metabolic correlates to HbS%. In **E**, overview of the top metabolic pathways that differ between high HbS and high HbA groups.

### Treatment with hydroxyurea had a minimal impact on RBC and plasma metabolism

As part of this study, a total of 306 subjects with SS genotype and HbA<20% were studied to understand the effects of hydroxyurea.^35^ Of these, 147 were in the SS group + hydroxyurea, compared to 159 untreated SS patients (**Figure 3.A**). We observed a positive association between hydroxyurea treatment and increments in HbF percentages (r < 0.5), the most significant observation in this analysis (p < 10^−17^ - **Figure 3.B**). However, hydroxyurea treatment was associated only with modest metabolic changes, namely significant increases in monounsaturated fatty acids (14:1 and 16:1), and decreases in RBC amino acids (asparagine, glycine, serine), carnosine, cytidine and purine oxidation products (allantoate – **Figure 3.C**), as confirmed via correlation of metabolomics data to HbF % (**Figure 3.D**).

**Figure 3.**
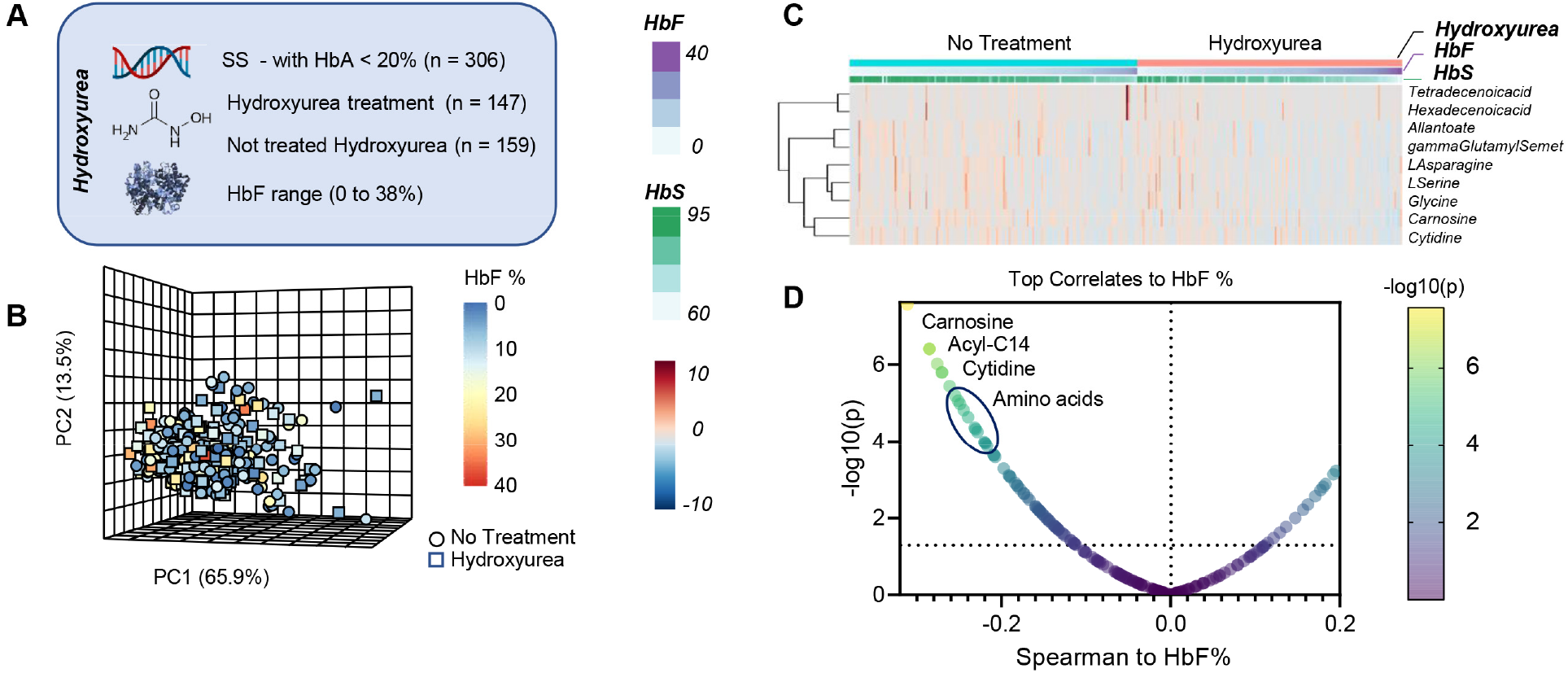
Metabolic correlates to Hydroxyurea treatment in RBCs from the WALK-PHaSST cohort. SS patients who were not recently transfused (HbA<20%; n=306) were tested as a function of treatment with hydroxyurea (n=147 vs n=159 untreated). In **B**, principal component analysis of the data from this cohort, color-coded based on HbF %. In **C**, heat map of the most significant metabolites affected by the treatment. In **D**, metabolic correlates to HbF%.

### Metabolic markers of hemolysis in the SS SCD population

SS subjects showed a significantly higher degree of hemolysis compared to other genotypes (p= 8.46 × 10^−26^) (**Supplementary Figure 5.A**). This finding validates our prior studies in two different cohorts of SCD patients, using plasma free hemoglobin and its association with clinical outcomes and risk allele loci in sickle cell patients.^14,35,46^ PCA of metabolomics data was performed in the SS population with HbA<20% (no HbC) – where each subject is color-coded as a function of the degree of hemolysis (**Figure 4.A**) – showed that PC1 (68.3% of the total variance) was significantly associated with hemolytic propensity. Hierarchical clustering analysis of these patients as a function of hemolysis highlighted progressive elevation in the levels of mono-, poly- and highly-unsaturated fatty acids (FA 18:1, 18:2, 18:3, 20:3, 20:4), acylcarnitines (C3, 12:1, 20:4), homocysteine, phosphocreatine, taurine and carnosine, urate – with concomitant decline in adenosine, PPP metabolite erythrose phosphate, serotonin and Acyl-C5 OH (**Figure 4.B**). These results are recapitulated and summarized in the smile plot in **Figure 4.C**, showing top correlates (Spearman) to the degree of hemolysis, including the tryptophan metabolite anthranilate.

**Figure 4.**
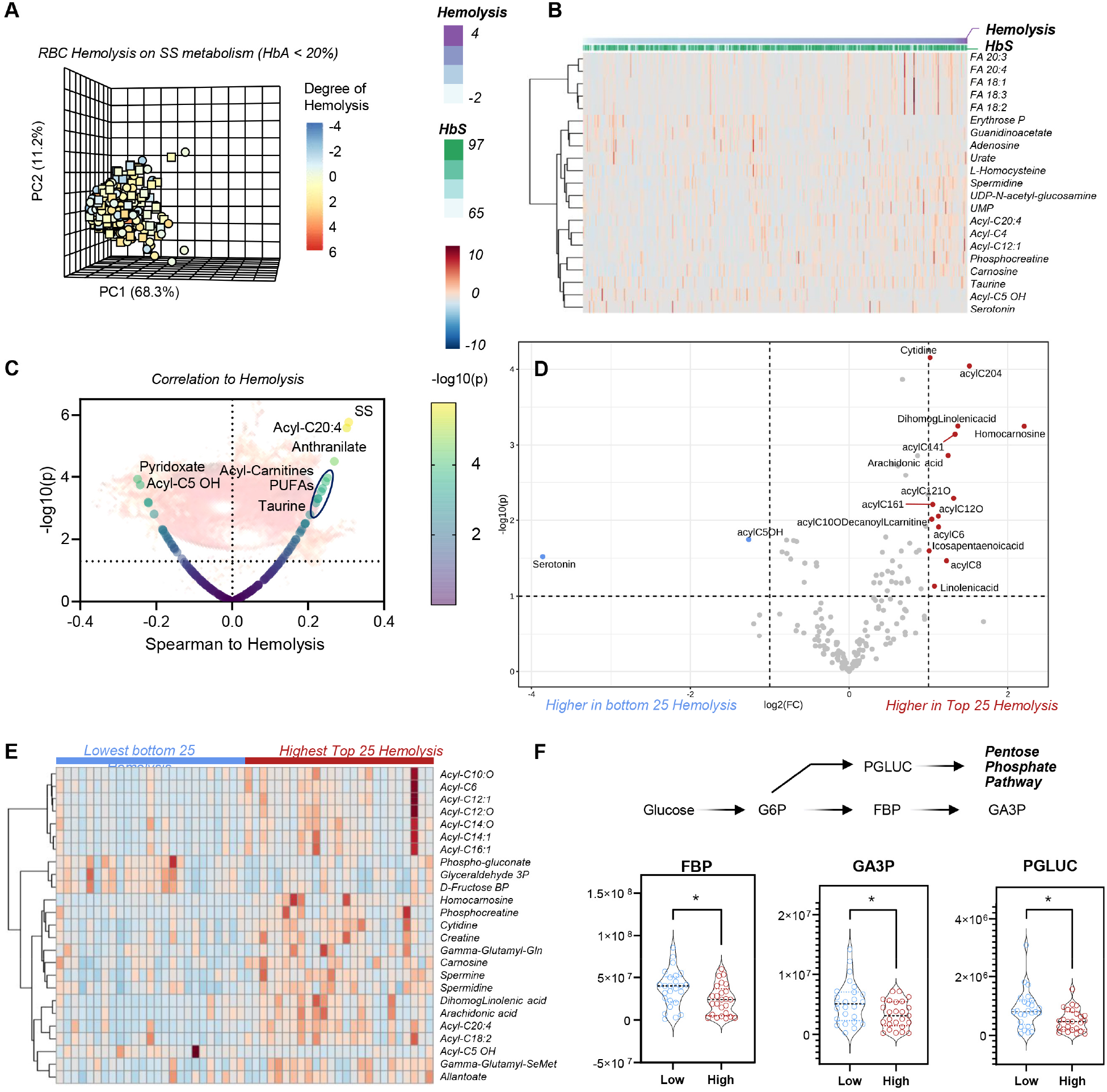
RBC metabolic markers of hemolysis in SS SCD patients who were not recently transfused (HbA <20%) In **A**, principal component analysis of the data from this cohort, color-coded based on the degree of hemolysis. In **B** and **C**, heat map and volcano plot of the metabolites most significantly associated with hemolytic propensity. In **D** and **E**, volcano plot and heat map ofthe most significant metabolic changes between the 25 patients with the highest and lowest degree of hemolysis, with a focus on metabolites of glycolysis and the pentose phosphate pathway (**F**).

We then repeated this analysis with either more or less stringent inclusion criteria. In the first analysis, we only focused on the SS patients, non-recently transfused, with the highest or lowest degree of hemolysis (n = 25 per arm – volcano plot and heat map following t-test in **Figure 4.D-E**). Elevation in acylcarnitines, declines in glycolysis (fructose bisphosphate, glyceraldehyde 3-phosphate) and the PPP (6-phosphogluconate) were associated with high hemolysis (**Figure 4.E-F**). Almost identical results were observed when analyses were repeated with no filter on genotypes, while still focusing on the 25 patients with extreme high or low hemolysis (**Supplementary Figure 5**). Consistent with a milder phenotype in the SC group, most patients in this analysis were SC patients, while higher hemolytic rate was associated with the SS group. This analysis, while confounded by the intrinsic differences between SS and SC RBCs, showed elevated S1P and depleted pyruvate in groups with highest hemolytic severity (**Supplementary Figure 5**).

### Metabolic correlates to Doppler-echocardiographic tricuspid regurgitant jet velocity (TRV) in the WALK-PHaSST cohort

TRV is associated with high pulmonary pressures and elevated prospective risk of death in adults with SCD. To identify RBC metabolic correlates to TRV elevation, we again focused on Hb SS subjects with HbA<20%. Results are shown in the form of a PCA, HCA or volcano plot in **Figure 5.A-C**, respectively. We identified a positive association with TRV for tryptophan metabolites (especially kynurenine, anthranilate, indoxyl, indole, indole 3-acetate), acylcarnitines (C5:1, 6 DC, 12:0, 20:4) - which were also associated with high hemolytic rate (**Figure 4.B**), arginine metabolites (citrulline, ornithine, creatinine and spermidine), succinate and urate (**Figure 5.B-C**). Increases in urate and depletion of 2-oxoglutarate were identified as the most significant positive and negative correlates to TRV when focused on subjects with extreme values (n=25 per group – **Figure 5.D**). Interestingly, a low plasma arginine is associated with high TRV, with the opposite observed in the RBC compartment. This divergent metabolic profile was observed for a number of metabolites, most notably arginine and S1P.

**Figure 5.**
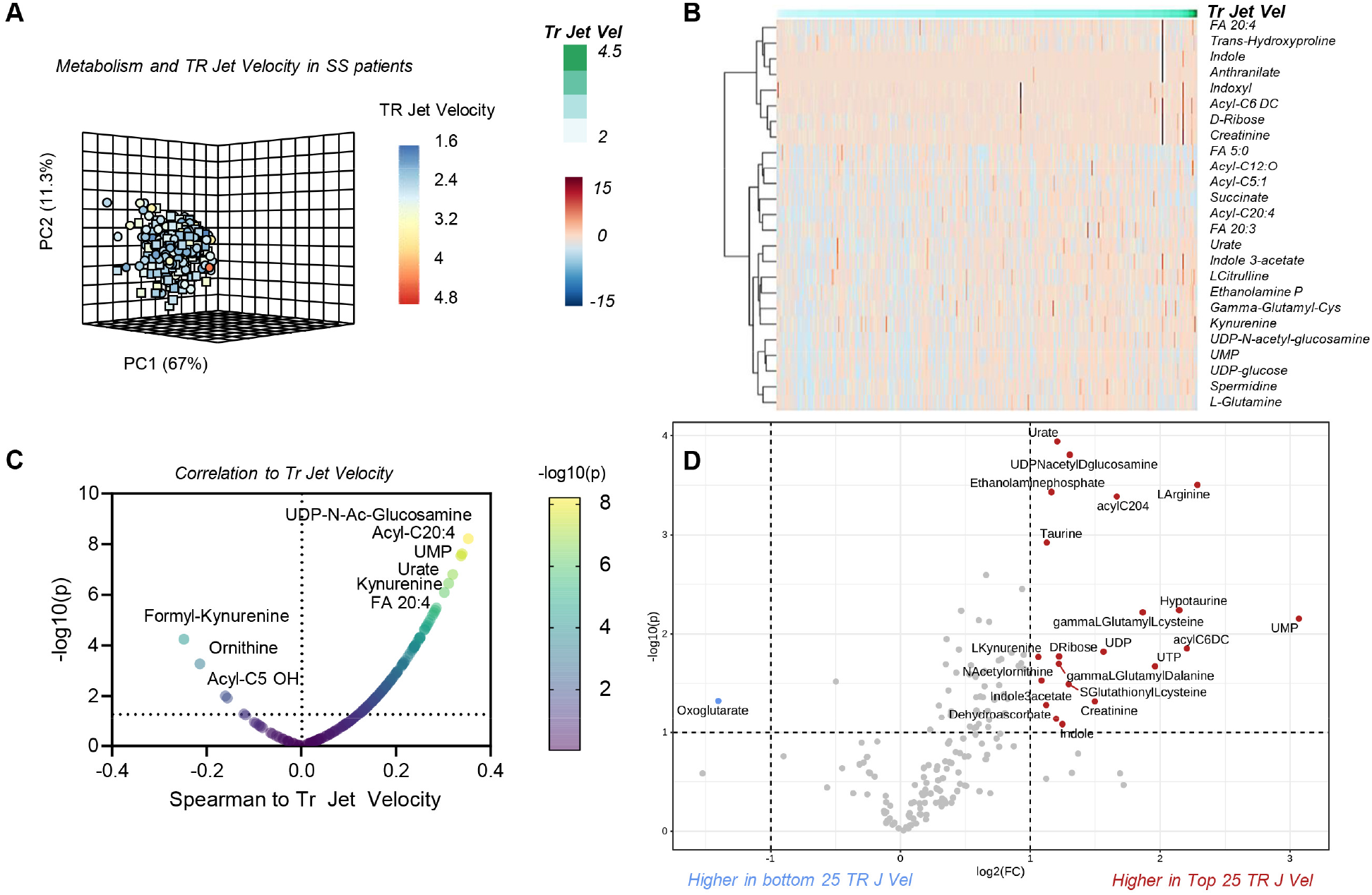
Metabolic markers of cardiovascular dysfunction as gleaned by TR_jet velocity in SS patients. In **A**, principal component analysis of the data from SS patients with no recent transfusion event (HbA<20%) in this cohort, color-coded based on the tricuspid regurgitation velocity (TRV). In **B** and **C**, heat map and volcano plot of the metabolites most significantly associated with TRV. In **D**, volcano plot and heat map of the most significant metabolic changes between the 25 patients with the highest and lowest degree of TRV.

### Metabolic correlates to renal dysfunction in the WALK-PHaSST cohort

Clinical and metabolic correlates to eGFR are shown in the PCA, HCA and volcano plot in **Figure 6.A-C**. Results show strong negative correlations (r < -0.5 Spearman, p < 10^−20^) for arginine metabolites (citrulline, creatinine), kynurenine, urate, succinate and acylcarnitines, and positive correlation with pyruvate, Acyl-C5 OH, ortho and diphosphate (**Figure 6.B-C**). These results were confirmed by more stringent analyses, focusing only on the extreme eGFR values (n=25 per arm) in SS patients (**Figure 6.D**). This analysis highlighted a positive association with eGFR for ATP, UTP, citrulline and eGFR, and a negative association with guanine, guanosine, GMP, AMP, dAMP, fructose bisphosphate and the Co-A precursor pantothenate (**Figure 6.D**). It is appreciated that changes in eGFR reflect chronic disease, and thus it is unclear if the associated changes in RBC metabolism reflect cause or effect of eGFR. Of note, eGFR declined with the age of the subject (**Figure 6.E**), with age and sex having a stronger effect on eGFR than genotype, when focusing on unadjusted data from the top 30 subjects with highest or lowest eGFR measurements (**Supplementary Figure 6**). Despite confounders, the most significant results from this less stringent analysis almost completely overlapped with the more stringent elaborations.

**Figure 6.**
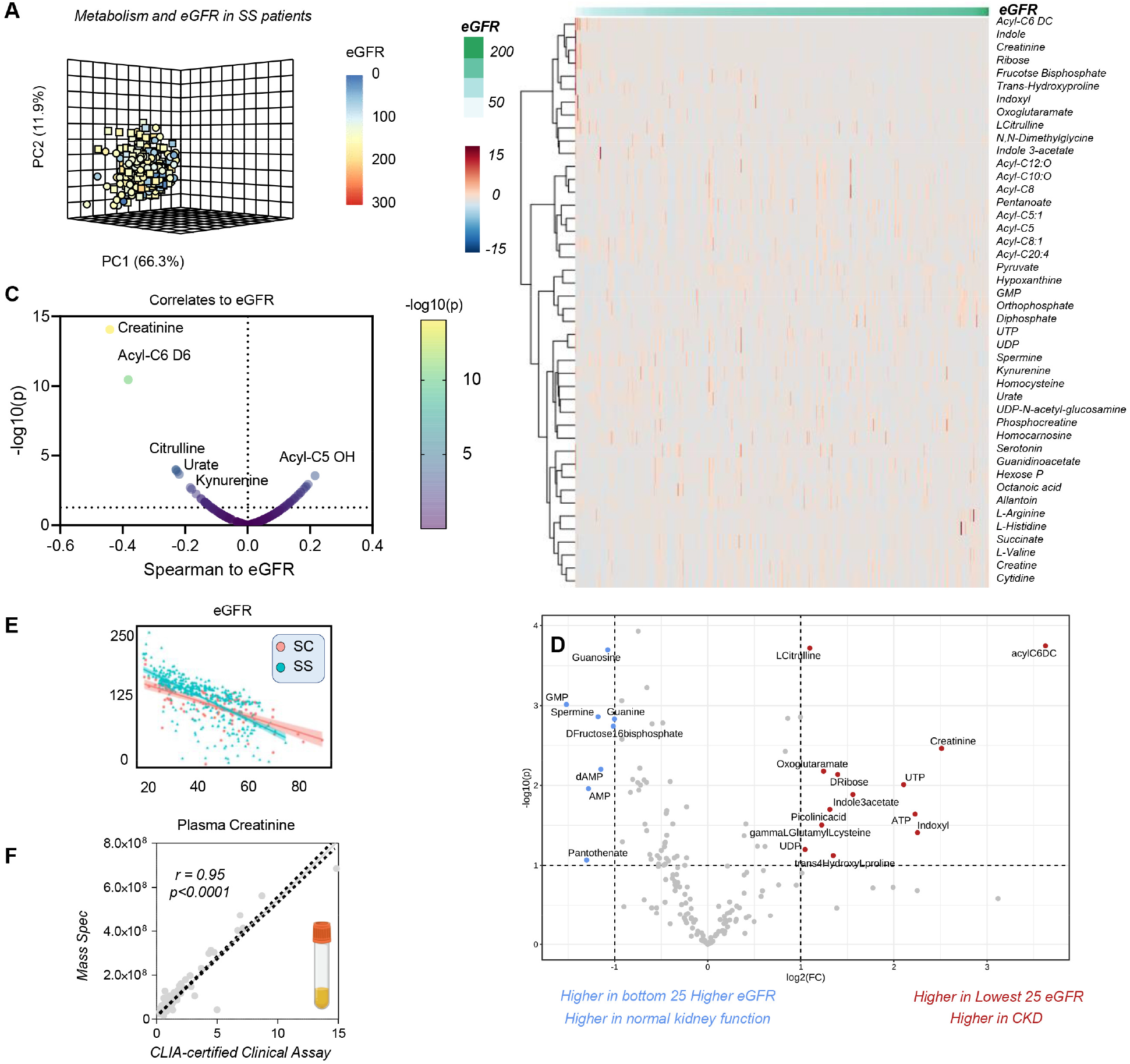
Metabolic markers of renal dysfunction as gleaned by eGFR and creatinine in SS patients. In **A**, principal component analysis of the data from this cohort, color-coded based on the eGFR. In **B** and **C**, heat map and volcano plot of the metabolites most significantly associated with eGFR. In **D**, volcano plot and heat map of the most significant metabolic changes between the 25 patients with the highest and lowest eGFR. In **E**, eGFR levels in SC and SS patients in the WALK PHaSST study as a function of age. In **F**, creatinine levels measured via CLIA certified clinical chemistry assay show a >0.95 correlation with the mass spectrometry data generated in this study.

As an internal cross-platform validation of our analytical approaches, clinical measurements of creatinine – a clinical marker of kidney dysfunction – were extremely reproducible between CLIA-certified clinical biochemistry assay and our mass spectrometry-based analysis (despite double randomization and blinding of the samples), confirming the quality of the data (**Figure 6.F**).

### RBC metabolic markers of mortality, time to event and hazard ratio in the WALK-PHaSST cohort

We next evaluated metabolic signatures of morbidity and mortality in the WALK-PHaSST cohort (**Figure 7**). Preliminary, crude unsupervised analyses were performed to discriminate metabolic markers of mortality in patients who deceased before the end of the study. Such analyses included PCA (**Figure 7.A**), hierarchical clustering (group averages of top 25 variables based on mortality are shown in **Figure 7.B**), and variable importance in projection (**Figure 7.C**). Duration of follow-up, a marker of mortality by design (**Figure 7.D**), and subject age were then used to calculate the time-to-event for patients who eventually deceased before the end of the study. Correlation to time-to-event suggests that the main variables associated with mortality in this population are CKD and cardiovascular risk, defined by a combination of TRV and GFR (**Figure 7.E**). Focusing on metabolic parameters only, biomarker analysis identified lactate as one of the top 10 predictors of time-to-death (ROC curve in **Figure 7.F**), along with creatinine (marker of CKD), purine deamination and oxidation products (urate, allantoin) and acylcarnitines. Notably, the heat map in **Figure 7.G** shows that time-to-death in SC and SS patients was accompanied by changes in three payoff phase glycolytic metabolites – phosphoglycerate isomers, phosphoenolpyruvate and pyruvate (**Figure 7.H**). To expand on these analyses and adjust for relevant covariates, we then performed a formal hazard ratio (HR) analysis, focusing on SS patients with lowest indication of recent transfusion (HbA < 10%). This analysis showed a significant association between acylcarnitine levels (especially acetyl- and Acyl-C5 OH pantothenate, glutamine and glutamyl-alanine, arachidonic acid (FA 20:4), creatinine and urate and decreased or increased hazard ratios (**Figure 7.G**).

**Figure 7.**
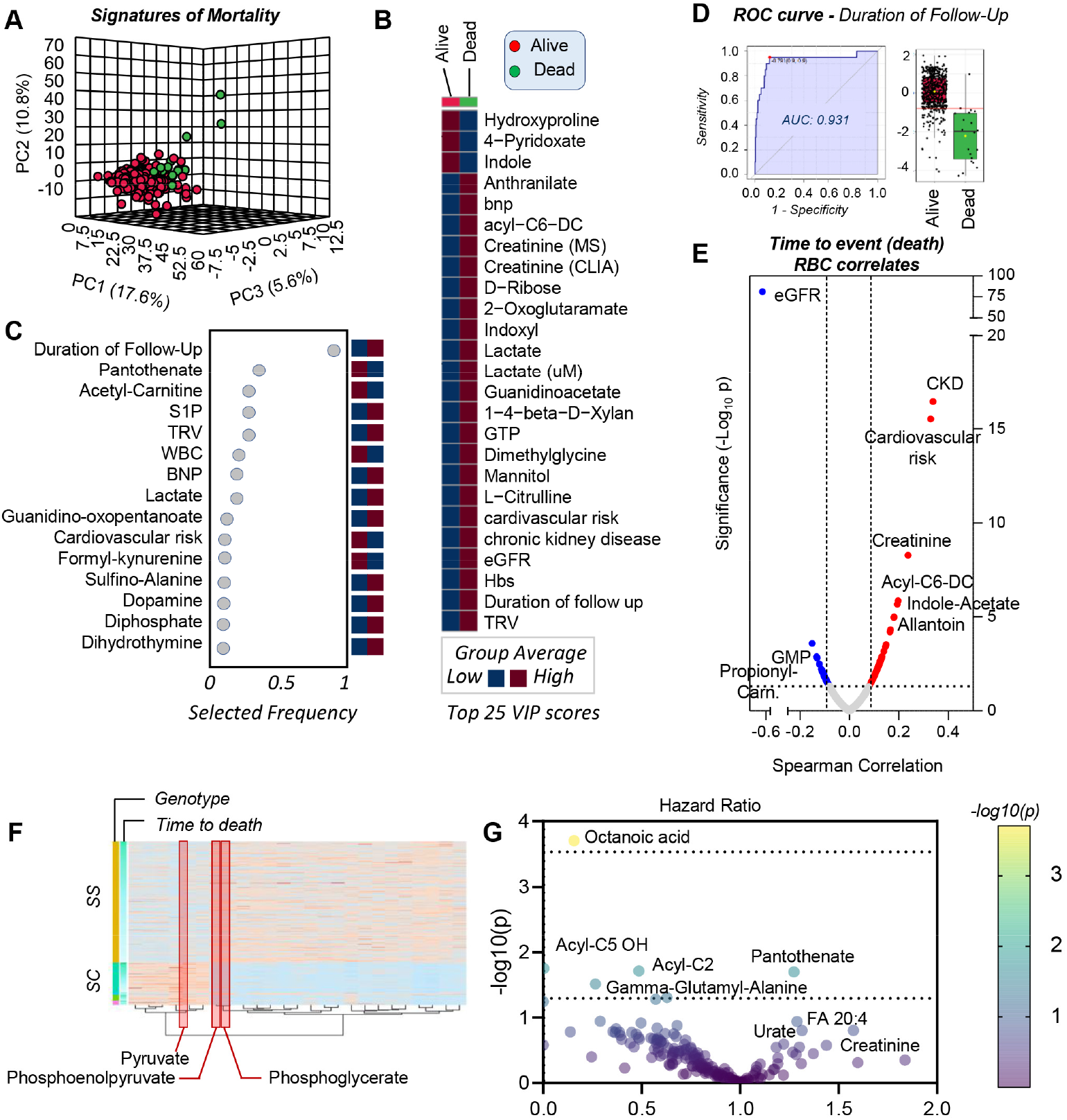
Clinical and metabolic signatures of mortality, time to event and hazard ratio in the WALK-PHaSST cohort. Unsupervised analyses, including PCA (green – dead; red – alive at the end of the study - **A**), hierarchical clustering (group averages of top 25 variables in **B**), variable importance in projection (**C**), receiver operating characteristic curves (ROC curve for the top marker – duration of follow-up – **D**) in the WALK-PHaSST cohort. Duration of follow-up and subject age were then used to calculate the time-to-death for patients who eventually deceased before the end of the study, which was then used to identify correlates to this variable (**E**). In the volcano plot in **E**, the X axis represents the linear correlation (r) values from Spearman correlationanalyses, and the Y axis indicates significance (-log10 p-values) for positive (red) or negative (blue) correlates. Biomarker analysis identified lactate as one of the top 10 predictors of time-to-death (**F**). In the heat map in **G**, results from the hazard ratio analysis of metabolomics data adjusted by relevant covariates.

### A free to access online portal for real time processing of metabolomics data in SCD

Given the wealth of clinical and RBC metabolomics data in this study, we created an online portal for real time data processing and figure generation. The portal, open and accessible at https://mirages.shinyapps.io/SickleCellDisease/ is meant to serve both as a repository and tool allowing for correlation analyses between metabolites and clinical covariates, with the opportunity to modify filters (e.g. age, sex, HbS, HbA, HbF) and correct for covariates of interest (**Supplementary Figure 7**).

## DISCUSSION

Here we performed an extensive metabolomics investigation of RBC samples from the WALK-PHaSST cohort, which was previously characterized at the clinical, physiological and genotypic levels.^23,33-38^ Importantly, because Walk-PHaSST performed hemoglobin electrophoresis at the time of baseline blood samples collection, we were able to compare RBC metabolomics between non-recently transfused Hb SS patients and patients who have >80% transfused Hb AA RBCs. Unexpectedly, despite a previously reported 44% decrease in hospitalization in SCD patients on hydroxyurea compared to placebo,^49^ effects reported here are limited to increases in HbF and decreases in WBC counts, with limited effects on RBC metabolism. On the other hand, a much stronger effect was noted in response to transfusion. Results show that HbA% is a strong a potential confounder in metabolomics studies in SCD patients undergoing exchange transfusion.^25^ Other than resulting in increases in the circulating levels of additive solution components^50^, elevated HbA levels upon transfusion corresponded not only to lower levels of RBC acylcarnitines, but also decreases in markers of hypoxia such as succinate and spermidine, immunomodulatory metabolites with pro-inflammatory, macrophage-activating activity,^51^ especially in the context of efferocytosis.^52^ Since these metabolites originate from or in response to dysfunctional mitochondria, it is interesting to note that transfusion does improve tissue oxygenation, thereby preventing mitochondrial uncoupling either in high-oxygen consuming tissues (including the kidney^53^ and heart) or in residual mitochondria that can be retained mature sickle RBCs.^54-56^

Excess (intra- and extra-vascular) RBC hemolysis is a driver of many comorbidities in SCD, as mediated by excess free iron,^57^ heme^58^ and nitric oxide scavenging hemoglobin.^59^ In keeping with previous findings,^35^ here we show that mortality and time-to-death in this cohort were strongly associated with the degree of renal and cardiovascular dysfunction, which in turn strongly correlated with the degree of hemolysis. Building on the observations above, we identified metabolic markers of hemolysis and the aforementioned comorbidities. One of the top biological correlates to these comorbidities was the age of the subject, which in turn was strongly positively associated with the RBC levels of creatinine (marker of kidney function^60^) and citrulline. These arginine^18^ metabolites have been previously associated with cardiovascular dysfunction and vaso-occlusive pain crisis,^24^ owing to the role as substrate for the synthesis of the vasodilator nitic oxide.^22,59,61^ Additional markers of disease severity included the elevation of intracellular acylcarnitines, a marker of RBC membrane lipid remodeling as a function of oxidant stress, through the activation of the Lands cycle.^27^ Interestingly, altered carnitine metabolism has been linked to increased RBC hemolysis following doxorubicin treatment - a common anthracycline in the treatment of hematological malignancies, and a trigger of renal and heart dysfunction in murine models.^62^ Carnitine supplementation may represent a therapeutic strategy to mitigate hemolysis and kidney dysfunction in SCD.^53^

Similarly, SCD disease severity corresponded to elevation of RBC S1P, an immunomodulatory metabolite that promotes T cell egression from the lymphatics, with a role in HbS stabilization and sickling in the hypoxic sickle RBC,^8^ neurocognitive impairment and cerebrovascular degeneration, vaso-occlusive crisis.^22^ S1P levels were elevated in RBCs from SS patients compared to SC subjects, suggestive of alterations in S1P synthesis via SphK1,^8,11^ or export via recently identified transporters. Notably, MFSD2B is an S1P transporter in the RBC,^63^ and SNPs of MFSD2B are associated with elevated hemolysis in the sickle and healthy RBC.^14^

Adenosine signaling dysregulation is a hallmark of hypoxia, SCD and related comorbidities.^9,64^ Here we report that purine deamination and oxidation are negatively impacted by HbS %, sex dimorphism (hypoxanthine-guanosine phosphoribosyl transferase is coded by an X-linked gene) and negatively correlates to hemolysis (especially urate, allantoin, xanthine and hypoxanthine), renal and cardiovascular function. Of note, murine models of kidney ischemia^65,66^ have shown that dysregulated purine metabolism (upon ATP breakdown and oxidation) is a hallmark of remote organ failure, especially in the heart, even a week after the ischemic event.^65^

Oxidant stress was exacerbated in the SS group – with depletion of GSH reservoirs (which emerged only upon correction for the impact of blood transfusion), activation of the PPP and accumulation of oxidized methionine and tryptophan metabolites, especially kynurenine. Elevation in the levels of kynurenine as a function of hemolysis and HbS is an interesting finding, in that kynurenine levels are a hallmark of cGAS-STING – Interferon – IDO1 axis activation in health and disease (e.g., COVID-19,^67^ Trisomy 21^68^), which in turn can be activated by circulating levels of nucleic levels, such as those observed in response to hemolysis of mitochondrial containing RBC and/or reticulocytes.^54^ Similarly, elevated levels of another class of tryptophan catabolites of bacterial origin, indoles, is indicative of potential microbiome dysbiosis in SCD,^69^ which has been recently associated with systems-wide pro-inflammatory events and, in particular, neurocognitive impairment.^70^ Oxidation of methionine and alterations of other one-carbon metabolites in one-carbon metabolism, like choline and serine as a function of disease severity could constrain methyl group availability for isoaspartyl-damage repair in SCD RBCs.^71^

Interestingly, we found that the SS genotype is associated with a metabolic blockade at a critical ATP-generating step of late glycolysis, which is catalyzed by redox sensitive PK.^72^ ATP is a key energy token to fuel RBC homeostatic processes, including degradation of oxidized components via ATP-dependent ubiquitinylation and proteasomal activity in SCD RBCs.^73^ PK oxidation state has been implicated in the etiology of asthma through regulation of pro-inflammatory cytokines (e.g., interleukin 1β)^74^ and dysregulated PK isoform splicing and activity is a hallmark of pulmonary hypertension,^75,76^ a common comorbidity in SCD. Recently, it has been shown that activation of PK with small molecule drugs has potent anti-sickling effects and improves RBC survival in a phase I clinical study^12^ and murine models of SCD.^77^ Our findings here are consistent with a decrease in PK activation in SCD patients with SS genotypes compared to clinically milder SC, suggesting that the former population would specifically benefit from PK targeting therapies.

## Acknowledgments

AD was supported by funds by the National Institute of General and Medical Sciences (RM1GM131968), and from the National Heart, Lung, and Blood Institute (R01HL146442, R01HL149714, R01HL148151, R01HL161004). The content is solely the responsibility of the authors and does not necessarily represent the official views of the National Institutes of Health.

## Authors’ contributions

MSN, YZ, VG, MTG designed and executed the WALK-PHASST study, collected and stored the samples, performed clinical measurements. FC, FG, JAR, AD performed metabolomics analyses (untargeted and targeted quantitative). KB and MG built the SCD metabolome portal. AD performed data analysis and prepared figures and tables. AD wrote the first draft of the manuscript, which was revised by all the other authors. All the authors contributed to finalizing the manuscript.

