## Supplementary Methods and Figures for "*In vivo* evaluation of the effect of sickle cell hemoglobin S, C and therapeutic transfusion on erythrocyte metabolism and cardiorenal dysfunction"

### SUPPLEMENTARY MATERIAL

**SUPPLEMENTARY DATA TABLE** *(PLEAE REFER TO "LEGEND" SHEET WITHIN THE FILE)* .......... **XLSX**

***Supplementary Materials and Methods - Extended***

***Ultra-High-Pressure Liquid Chromatography-Mass Spectrometry (MS) metabolomics:*** Frozen RBC aliquots (50 µL) were thawed on ice then extracted 1:10, in ice cold extraction solution (methanol:acetonitrile:water 5:3:2 *v/v/v*). Samples were vortexed for 30 min at 4℃ and insoluble material pelleted via centrifugation at 15,000 *g* for 15 min under refrigerated conditions, as described.[1,2] Analyses were performed using a Vanquish UHPLC coupled online to a Q Exactive mass spectrometer (Thermo Fisher, Bremen, Germany). Samples were resolved as described,[3,4] over a Kinetex C18 column (2.1x150 mm, 1.7 µm; Phenomenex, Torrance, CA, USA) at 45°C. A volume of 10 ul of sample extracts was injected into the UHPLC-MS. Each sample was injected with two different chromatographic and MS conditions as follows: 1) using a 5 minute gradient at 450 µL/minute from 5-95% B (A: water/0.1% formic acid; B:acetonitrile/0.1% formic acid) and the MS was operated in positive mode and 2) using a 5 minute gradient at 450 µL/minute from 5-95% B (A: 5% acetonitrile, 95%water/1 mM ammonium acetate; B:95%acetonitrile/5% water, 1 mM ammonium acetate) and the MS was operated in negative ion mode. The UHPLC system was coupled online with a Q Exactive scanning in Full MS mode at 70,000 resolution in the 60-900 m/z range, 4 kV spray voltage, 15 sheath gas and 5 auxiliary gas. These chromatographic and MS conditions were applied for both relative and targeted quantitative metabolomics measurements, with the differences that for the latter targeted quantitative post hoc analyses were performed on the basis of the stable isotope-labeled internal standards used as a reference quantitative measurement, as detailed below.

***Quality control and data processing:*** Calibration was performed prior to analysis using the Pierce™ Positive and Negative Ion Calibration Solutions (Thermo Fisher Scientific). Acquired data was then converted from raw to .mzXML file format using RawConverter. Samples were analyzed in randomized order with a technical mixture injected every 15 samples to qualify instrument performance and ensure technical coefficients of variations (standard deviation divided by the mean) below 20%. Metabolite assignments, isotopologue distributions, and quantification of stable isotope-labeled internal standards were performed using MAVEN (Princeton, NJ, USA), as described.[5]

***Statistical analyses:*** Data analysis was performed through the auxilium of the software MAVEN. Graphs and statistical analyses (either two-way ANOVA or repeated measures ANOVA) were prepared with GraphPad Prism 9.0 (GraphPad Software, Inc, La Jolla, CA), GENE E (Broad Institute, Cambridge, MA, USA), Multivariate analyses, including principal component analyses (PCA), hierarchical clustering analyses, two-way ANOVAs, correlation analyses (Spearman) and calculation of receiver operating characteristic (ROC) curves were performed through the software MetaboAnalyst 5.0.[6] For survival analysis, we used time to right censorship (including death or last follow-up) as time to event and vital status (death or alive) was the studied outcome. PCA was used to derive a hemolytic component from lactate dehydrogenase, aspartate aminotransferase total bilirubin and reticulocyte percent.[7-9] We applied Cox proportional hazard models to calculate the hazard ratios and P value for each metabolite. and Cox analysis and adjusted regressions of metabolites to clinical outcomes were performed in R (R Core Team (2022), https://www.r-project.org/).

***Sickle Cell Disease ShinyApp portal*** A portal for online sharing of all the data generated in this study, in like-wise fashion to our recent COVID-Ome Explorer portal.[10] After data curation and quality control, each of the datasets (RBC metabolomics, clinical data) was linked at the sample level with a unique identifier, enabling cross-referencing among datasets. Then, each of the datasets was imported into applications developed using R, R Studio, and the R-based web application framework Shiny. All code required to run the COVIDome Explorer applications can be found at https://github.com/cusom/CUSOM.COVIDome.Shiny-Apps (Zenodo https://doi.org/10.5281/zenodo.5081091) and https://github.com/cusom/CUSOM.ShinyHelpers (Zenodo https://doi.org/10.5281/zenodo.5081093).

#### Supplementary Figures

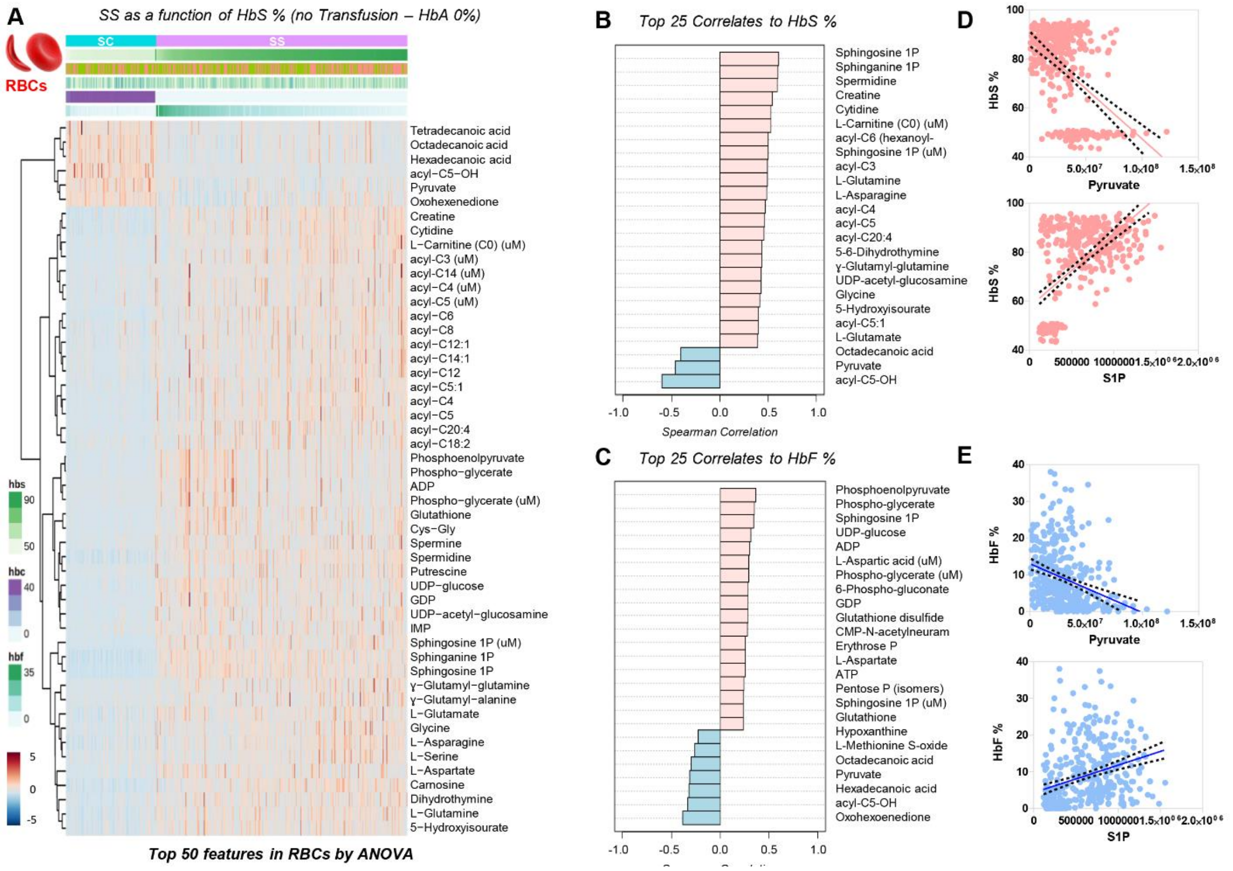

**Supplementary Figure 1** ***Analysis of metabolomics data in SS and SC patients from the WALK-PHaSST cohort as a function of HbS in the absence of transfusion*** *(HbA = 0%; SS n = 275; SC n = 99)*. Results are shown in the form of heat maps and hierarchical clustering of the top 50 metabolites in RBCs (A)and plasma (B). Top 25 metabolic correlates to HbS % and HbF% are shown in C and D, respectively. Scatter plots for top correlates of selected metabolites are shown in E and F.

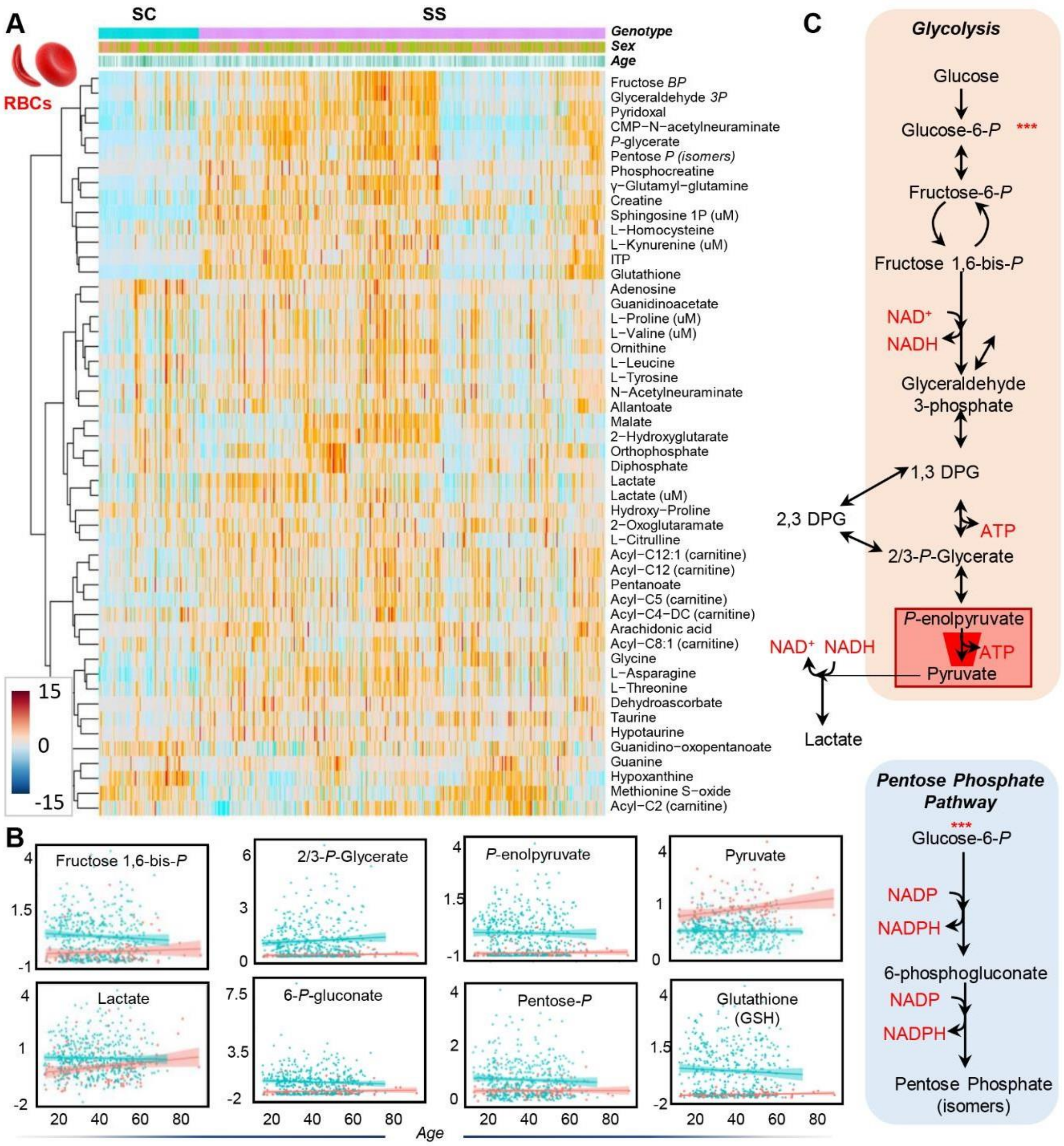

**Supplementary Figure 2** ***Metabolic differences between RBCs from subjects with SC and SS genotypes*** In **A**, top 50 RBC metabolites as a function of SC (n =107) vs SS (n=429) genotypes across all WALK-PHaSST samples. In **B**, an overview of glycolysis and the pentose phosphate pathway and in **C,** line plots for metabolites in these pathways that vary significantly between SC (SC) and SS subjects (green), as a function of subject age (X axis). Opposite trends between SC and SS for phosphoenolpyruvate (substrate, higher in SS) and pyruvate (product, along with ATP), higher in SC (**B**). Data are suggestive of a metabolic bottleneck in the ATP-generating glycolytic step catalyzed by the enzyme pyruvate kinase (**C**).

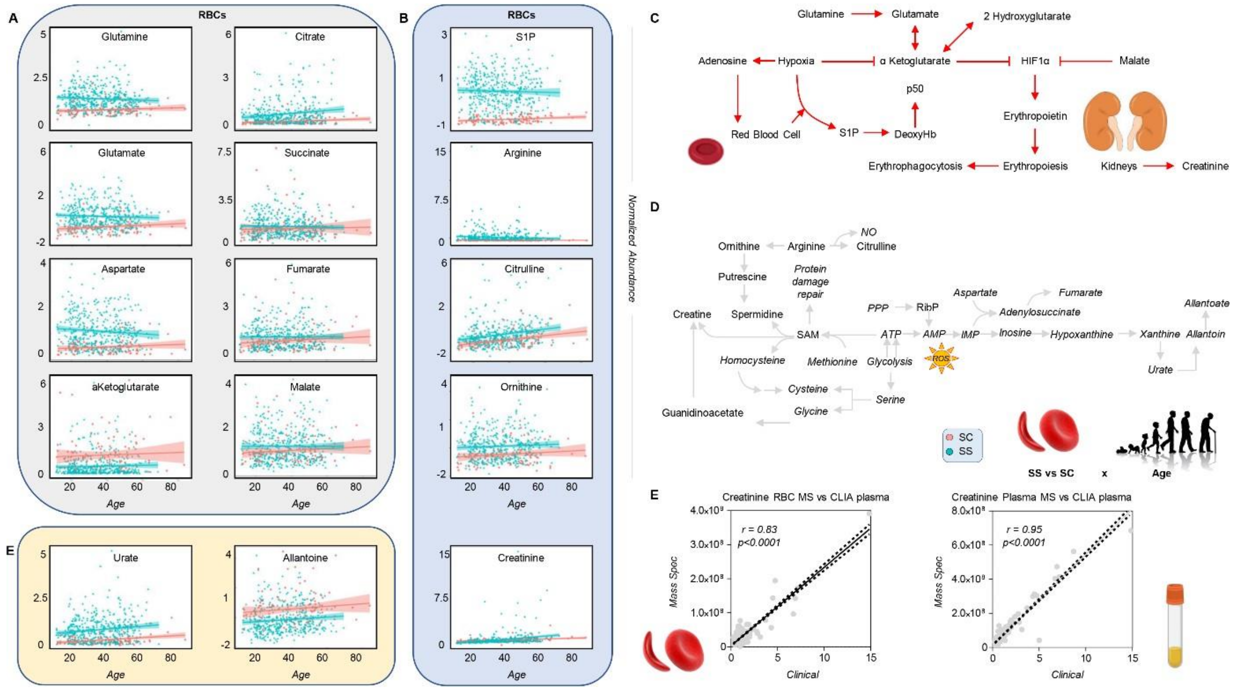

**Supplementary Figure 3** ***Metabolic differences in carboxylic acids, purine and arginine metabolism between RBCs from sickle cell disease patients with SC or SS genotypes*** In **A,** line plots for metabolites in carboxylic acids, purine and arginine metabolism that vary significantly between SC (SC; n=107) and SS subjects (green; n=429), as a function of subject age (X axis). In **C** and **D**, an overview of these metabolic pathways in mature RBCs. In **E**, correlation of creatinine levels detected by mass spectrometry in RBC (left) and plasma (right) compared against plasma measurements of creatinine, as determined via clinical assays.

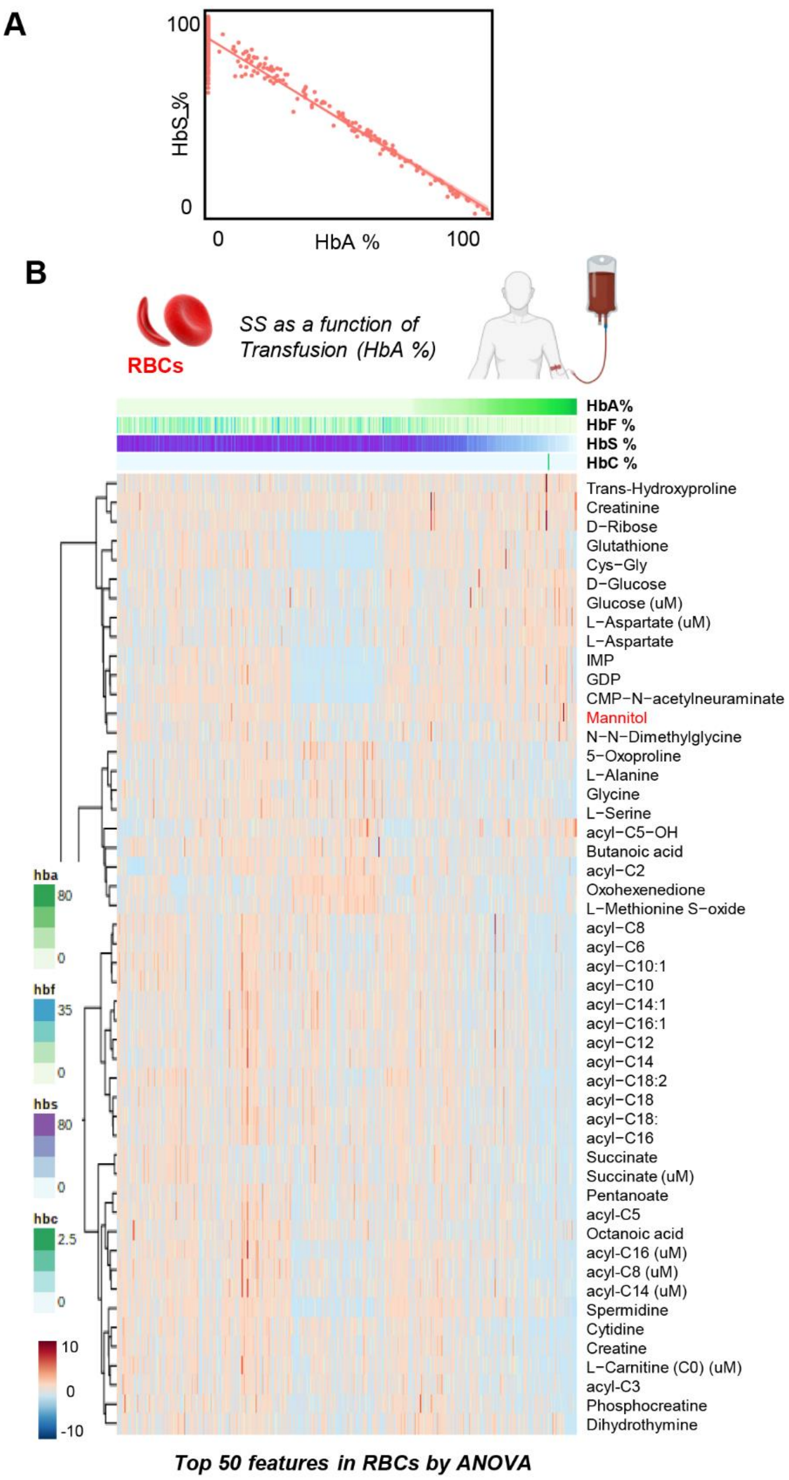

**Supplementary Figure 4** ***Impact of transfusion on the RBC metabolome in sickle cell patients as a function ofhemoglobin A %*** SCD patients in the WALK-PHaSST cohort (SS genotype – n=429, without excluding patients on hydroxyurea) were sorted as a function of HbA %, an indicator of recent transfusion. The top 50 significant RBC metabolic correlates to transfusion are shown in the heat map in A. Meta-data from this analysis clearly show a negative correlation between HbA% and HbS% (A), as expected. In B, thewe show the top 50 metabolites impacted by transfusion as a function of HbA.

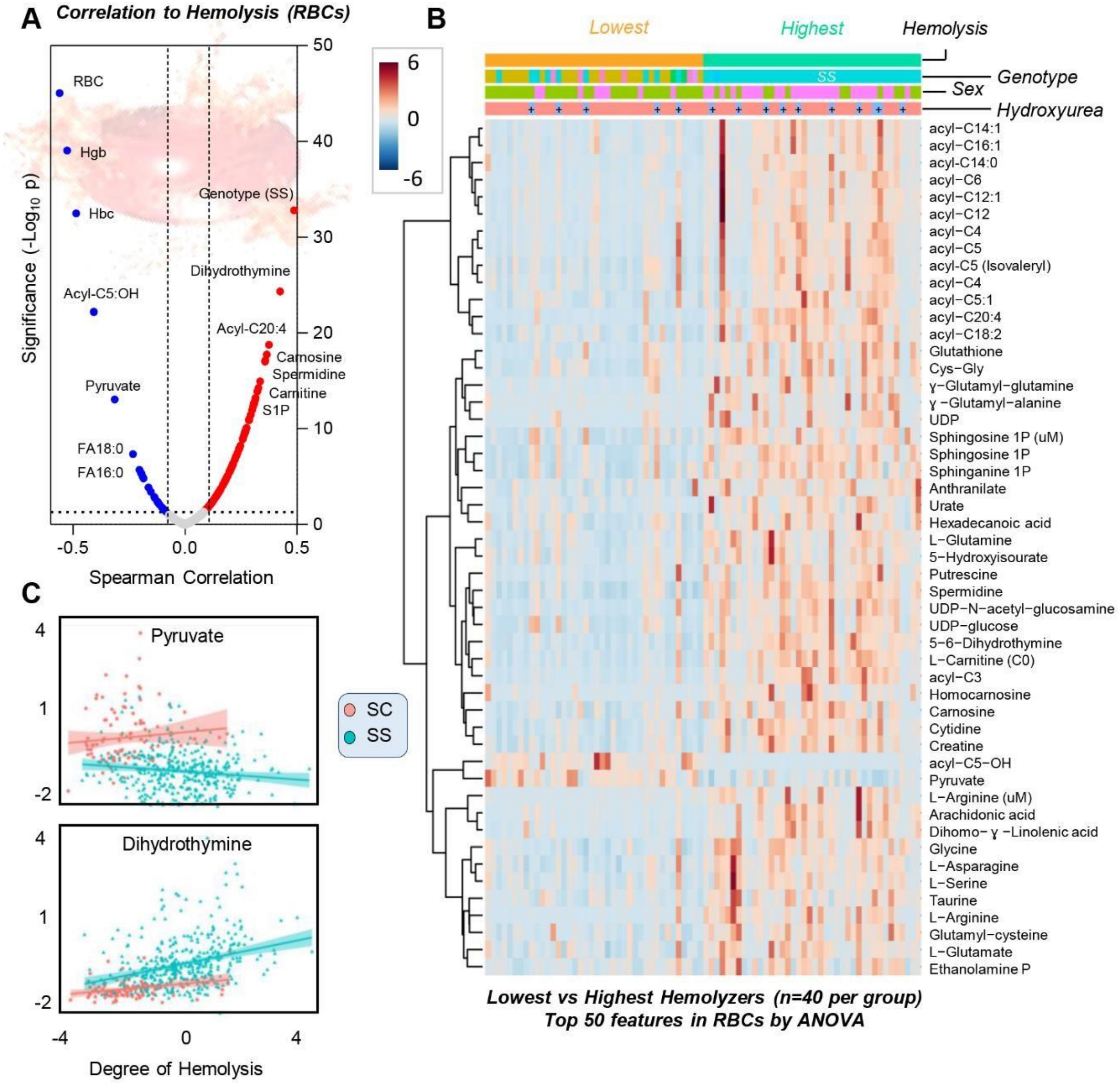

**Supplementary Figure 5** ***Metabolic markers of hemolysis in RBCs from the WALK-PHaSST cohort*** In the volcano plot in **A**, the X axis represents the linear correlation (r) values from Spearman correlation analyses of RBC metabolites and the degree of hemolysis, and the Y axis indicates significance (-log10 p-values) for positive (red) or negative (blue) correlates. In **B**, top 50 RBC metabolites as a function of lowest and highest hemolysis (n =40 per group) across all WALK-PHaSST samples. In **C**, two of the top metabolic correlates are shown, RBC pyruvate and dihydrothymine (X axis is the degree of hemolysis and Y axis is normalized abundance of the metabolite) for sickle cell SC genotype (red line) vs SS genotype (green line).

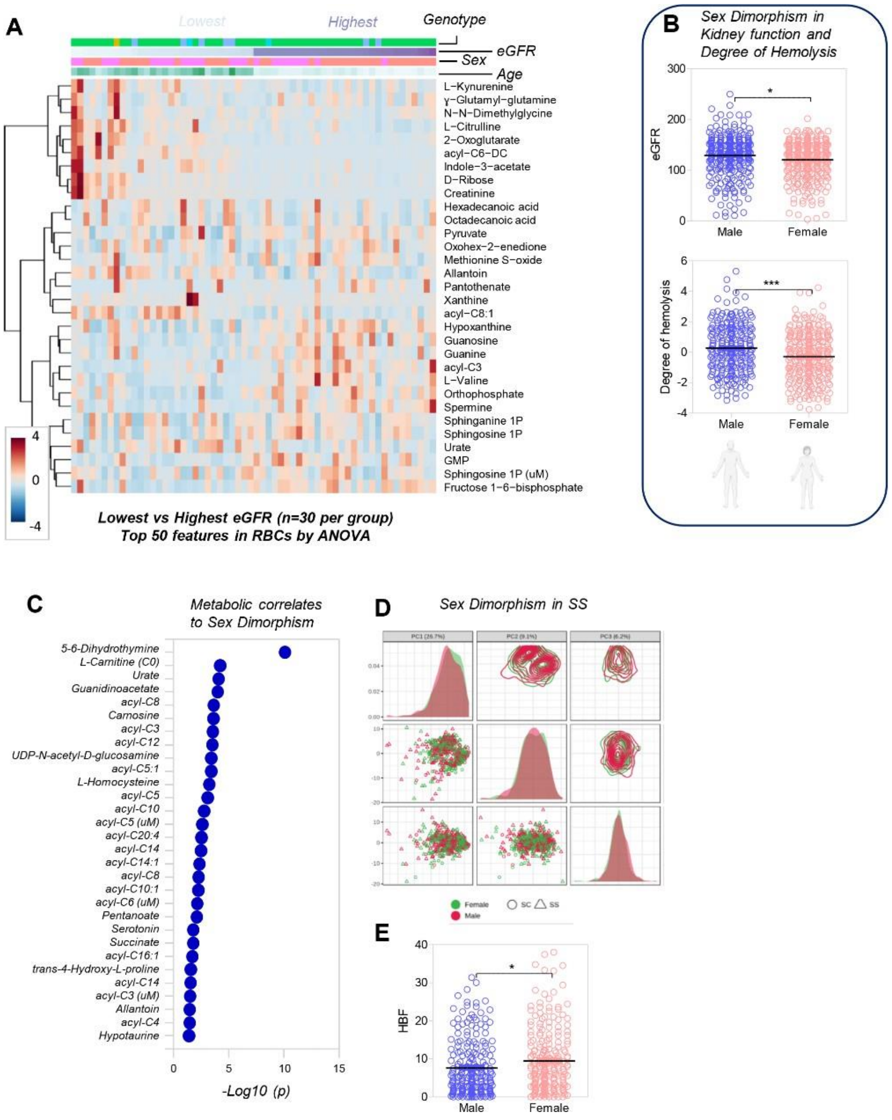

**Supplementary Figure 6** ***Impact of sex dimorphism in clinical and metabolic covariates in the WALK-PHaSST cohort*** Meta-analysis of RBC metabolomics data from subjects with the 30 lowest and highest eGFR confirmed that women were over-represented in the group with the lowest eGFR (n=30) and mostly correlated with higher levels of RBC kynurenine and other tryptophan metabolites of bacterial origin (indoles), carnitines (A). In this cohort, women showed significantly lower estimated glomerular filtration rates (eGFR – p<0.05) and degree of hemolysis (p<0.001 - **B**). The top 25 metabolites impacted by sex dimorphism are shown in panel **C**, as determined by variance importance in projection from principal component analyses – with multiple acyl-carnitines in this list (**E**). In **D**, women had significantly higher levels of hemoglobin F (HbF).

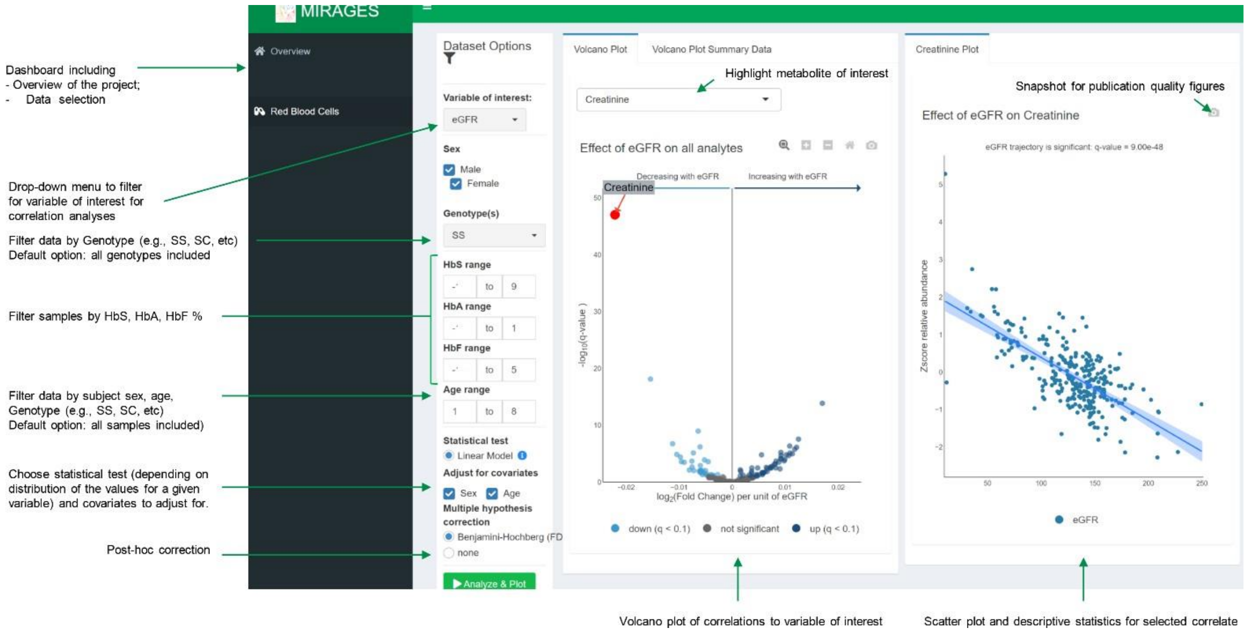

**Supplementary Figure 7** ***Overview of the sickle cell metabolome portal*** For this particular analysis, we focused on RBC metabolic correlates to eGFR, after selecting for SS genotype only, for any subject with HbA < 10%. The portal is freely accessible for interactive exploration and elaboration of the data presented in this study at https://mirages.shinyapps.io/SCD/
